# BRG1 defines a genomic subset of inflammatory genes transcriptionally controlled by the glucocorticoid receptor

**DOI:** 10.1101/2021.12.13.472398

**Authors:** Aikaterini Mechtidou, Franziska Greulich, Benjamin A. Strickland, Céline Jouffe, Filippo M. Cernilogar, Gunnar Schotta, N. Henriette Uhlenhaut

## Abstract

Glucocorticoids (such as Dexamethasone) are commonly used immunomodulatory drugs with potent anti-inflammatory effects, whose mechanisms of action remain incompletely understood. They bind to the Glucocorticoid Receptor (GR), a nuclear hormone receptor that acts as a transcription factor to directly control the expression of inflammatory genes. To elucidate the complex molecular mechanisms employed by GR during the suppression of innate immune responses, we have performed proteomics, ChIP-seq, ATAC-seq, RNA-seq and bioinformatics together with genetic and pharmacological loss of function studies in primary mouse macrophages. We found that GR interacts with the ATP-dependent SWI/SNF chromatin remodeling complex to regulate a specific subset of target genes. Here we show that the central catalytic subunit BRG1 is required not only for the transcriptional activation of classical GR target genes such as *Fkbp5* or *Klf9*, but also for the transcriptional repression of cytokines and chemokines such as *Ccl2, Cxcl10* or *Il1a*. We demonstrate that loss of BRG1 activity leads to reduced histone deacetylase (HDAC) function, and consequently increased histone acetylation, at these repressive GR binding sites. Altogether, our findings suggest that GR interacts with BRG1 to assemble a functional co-repressor complex at a defined fraction of macrophage *cis*-regulatory elements. These results may indicate additional non-classical, remodeling-independent functions of the SWI/SNF complex and may have implications for the development of future immunomodulatory therapies.

**GRAPHICAL ABSTRACT:** Graphical Abstract.
In macrophages (mΦ) responding to bacterial LPS and Dexamethasone, the Glucocorticoid Receptor (GR) activates target genes like *Klf9* or *Fkbp5* via interaction with the BRG1-containing SWI/SNF complex, chromatin remodeling and Mediator recruitment. At the same time, GR represses the expression of inflammatory cytokines and chemokines such as *Ccl2, Cxcl10, Il1a etc*. by assembling a BRG1-containing co-repressor complex and de-acetylating surrounding histone tails. Loss of BRG1 activity affects both the transcriptional activation and repression of a subset of myeloid GR target genes via distinct mechanisms. (iTF: inflammatory transcription factor; Ac: histone acetylation) (Created with BioRender.com.)

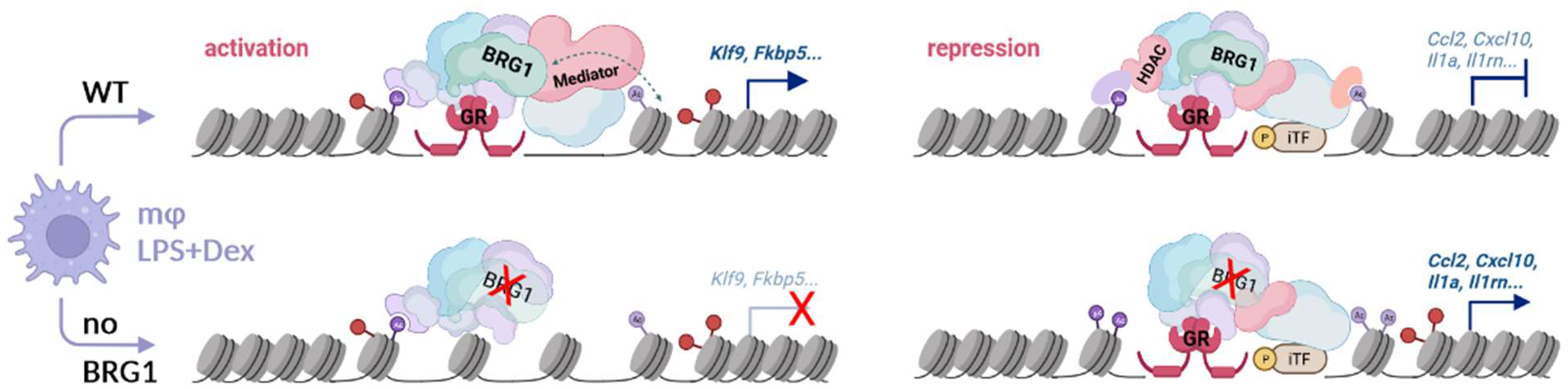

## INTRODUCTION

The Glucocorticoid Receptor (GR, encoded by the *Nr3c1* gene) is an important immunomodulatory drug target and a prominent physiological regulator. It belongs to the nuclear receptor family of ligand gated transcription factors, whose clinical relevance is underscored by its life-saving effects in COVID-19 patients (Group et al. 2021). Upon binding to its ligands such as Dexamethasone, GR translocates to the nucleus to either activate or repress target gene transcription. The exact mechanisms specifying positive versus negative regulation and the composition of coregulatory complexes assembled on target promoters or enhancers are inherently complex and pose many open questions (Escoter-Torres et al. 2019). Several studies have suggested that *cis*-regulatory element recognition and binding by GR is predetermined by each cell type’s specific chromatin landscape, which is established by pioneer factors like PU.1, AP-1 or C/EBP, and which shapes the GR cistrome (Biddie et al. 2011; John et al. 2011; Greulich et al. 2016).

In that respect, chromatin remodeling is both an essential prerequisite as well as a central component of GR-mediated transcriptional regulation. Assembly of the SWI/SNF (SWItch/Sucrose-NonFermentable) remodeling complex and its interaction with GR have been shown to enhance the transcriptional hormone response. BRG1 (SMARCA4), the central ATPase of the SWI/SNF complex, is required for proper and robust GR-regulated gene activation (Fryer and Archer 1998). Both structural models and biochemical experiments indicate that SMARCD1 (BAF60A), SMARCC1 (BAF155), SMARCE1 (BAF57) and ARID1A (BAF250) components engage in protein-protein interactions between GR and the SWI/SNF complex (Hsiao et al. 2003; Muratcioglu et al. 2015). Gene activation of various nuclear receptors, including GR, has been reported to broadly require the cooperation of this well-studied chromatin remodeling complex. In this context, BRG1 both precedes GR chromatin occupancy, by establishing pioneer factor recruitment to create accessible GR DNA binding sites, and also serves subsequently as a coactivator and remodeler required for GR-induced DNA accessibility and transcription (Trotter and Archer 2004; Trotter et al. 2015; Hoffman et al. 2018).

Regarding its clinical use, the direct transcriptional repression of pro-inflammatory cytokines and chemokines by GR is thought to underlie a major part of its immunomodulatory potency (Escoter-Torres et al. 2019). Indeed, gene repression was partially affected in 3134 cells expressing a dominant negative BRG1, and individual glucocorticoid-induced BRG1-dependent DNAse hypersensitivities were described. *John et al*. suggested an important role of chromatin remodeling in GR-mediated repression, based on the detection of numerous transition events linked to repressed loci (John et al. 2008). Furthermore, BRG1 was found to be required together with HDAC2 for histone de-acetylation and repression of the human *POMC* promoter, a well-known negative GR target (Bilodeau et al. 2006).

Finally, genome wide studies during the past decade have revealed that GR binding sites are not only cell type-, signal- and time point-specific, but that given GR cistromes are far from uniform, and can be divided into distinct subsets or particular classes of target loci. We therefore hypothesized that BRG1-containing remodeling complexes may mediate significant fractions of anti-inflammatory glucocorticoid actions. Here we chose primary bone marrow derived murine macrophages, which are important cellular mediators of the innate immune response, as a model to study the GR-mediated repression of inflammatory genes (Uhlenhaut et al. 2013; Greulich et al. 2021b). We performed ChIP-seq and ATAC-seq in lipopolysaccharide-activated macrophages to functionally characterize the role of BRG1 (SMARCA4) for a subset of GR target genes. Our data suggest that the catalytic activity of the SWI/SNF complex is not only involved in the activation of classical GR target genes (such as *Klf9* or *Fkbp5*), but also in the transcriptional repression of pro-inflammatory cytokines, chemokines and interleukins (such as *Ccl2, Cxcl10* or *Il1a*).

## RESULTS

### GR and BRG1 co-occupy macrophage *cis*-regulatory loci

In order to chart the composition of the transcriptional complexes assembled by GR during the regulation of innate immune responses, we performed protein-protein interactome mapping by ChIP-MS for GR in primary murine bone marrow derived macrophages activated with the TLR4 ligand lipopolysaccharide (LPS) and treated with the GR ligand Dexamethasone (Dex) (Greulich et al. 2021b). In addition to various known co-regulators and to novel interaction partners such as the COMPASS complex, we found several components of the SWI/SNF complex significantly enriched together with GR (**Fig. 1A**). For example, we detected SMARCD2 (BAF60B), SMARCE1 (BAF57), SMARCC2 (BAF170) and ARID1A (BAF250) peptides in our IP dataset. To confirm these putative interactions between GR and SWI/SNF subunits in activated macrophages, we then carried out endogenous Co-IPs in the RAW264.7 myeloid cell line, in the presence of LPS and Dex. Indeed, we were able to detect GR together with SMARCD1 (BAF60A), SMARCE1 (BAF57) and SMARCA4 (BRG1) by Western Blotting (**Fig 1B)**.

**Figure 1:**
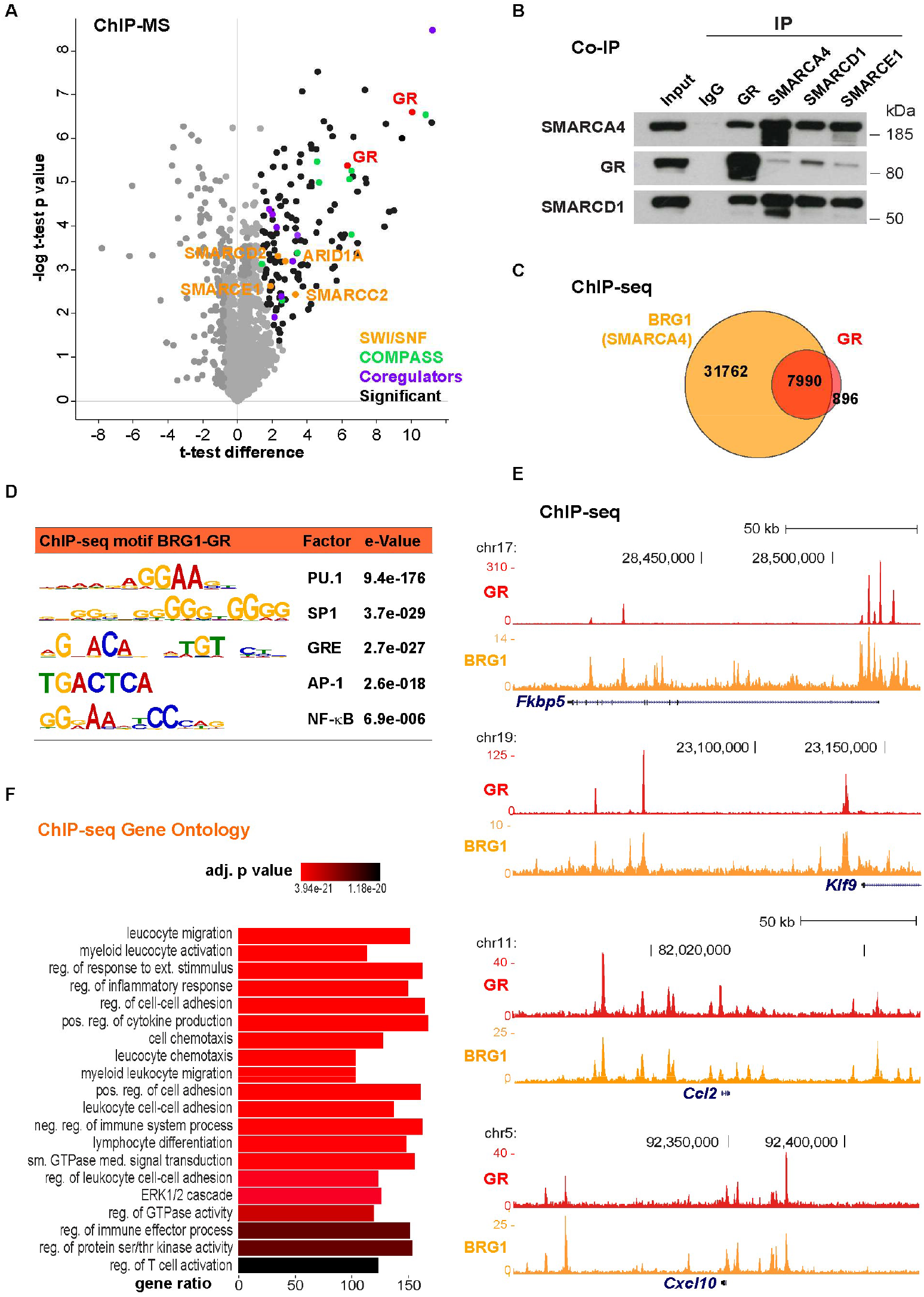
GR interacts with the SWI/SNF complex in activated macrophages. (A) ChIP-MS GR interactome in primary macrophages treated with Dexamethasone (Dex) and lipopolysaccharide (LPS) (Greulich et al. 2021b). Colored dots show interactors significantly enriched in a GR pulldown over non-specific isotype-matched IgG, functionally annotated (1.5fold, p<0.05). (B) Western blot of endogenous Co-IPs in RAW 264.7 cells treated with LPS and Dex. (C) Venn diagram of reproducible GR and BRG1 ChIP-seq peaks overlapping in primary macrophages treated with LPS and Dex (n=2). (D) Motif enrichment analysis for the 7,990 common GR and BRG1 ChIP-seq peaks. (E) Representative genome browser tracks of GR and BRG1 ChIP-seq signals, showing means from two replicates. (F) Functional annotation of the 7,990 common GR-BRG1 sites, assigned to the nearest gene.

Similarly, when we compared our macrophage interactome with data from livers and Dex-treated mouse embryonic fibroblasts (MEFs, activated by LPS), we also found the SWI/SNF subunits to be enriched (Fig. S1A) (Quagliarini et al. 2019; Escoter-Torres et al. 2020). Therefore, we conclude that GR robustly interacts with the SWI/SNF chromatin remodeling complex across tissues and cell types.

To further investigate potential functional relationships between GR and the SWI/SNF remodeling complex, we next performed ChIP-seq for both GR and the core ATPase subunit BRG1 (SMARCA4) in primary murine macrophages. (Since the other catalytic SWI/SNF component, SMARCA2 (BRM), was transcriptionally downregulated upon Dex stimulation, we focused only on BRG1 (Fig. S1B)). As shown in **Fig. 1C**, almost all GR binding sites mapped in response to LPS and Dex, also showed co-occupancy of BRG1 (about 90%). As expected, we also detected many additional BRG1 binding sites throughout the genome, not overlapping with GR, which represent the central, essential functions of the SWI/SNF complex within the macrophage chromatin landscape (Chen et al. 2020). Bioinformatic motif analyses of those ∼8,000 common GR-BRG1-bound ChIP sequences revealed the GR consensus motif (GRE) as significantly enriched, together with the known macrophage pioneer factor PU.1 and the inflammatory mediators AP-1 and NF-κB (**Fig. 1D**, Fig. S1C), validating our data sets (Uhlenhaut et al. 2013). For instance, we observed BRG1 binding to GR target sites such as the *Fkbp5, Klf9, Ccl2, Cxcl10, Il1a and Il1rn* loci (**Fig. 1E**, Fig. S1D). *Fkbp5* and *Klf9* are two typical examples of positive GR targets *induced* by Dex, while *Cxcl10, Ccl2, Il1a, and Il1rn*, are representative cases of negative GR target genes *repressed* in response to ligand (Uhlenhaut et al. 2013; Escoter-Torres et al. 2020; Greulich et al. 2021b). In addition to these exemplary cytokines, the functional annotation of the ∼8,000 common GR-BRG1 target sites, based on the nearest gene, include many genes involved in inflammation, immune responses, myeloid migration and inflammatory signaling cascades (**Fig. 1F**).

Altogether, our immunoprecipitation studies in macrophages show that the central SWI/SNF component BRG1 co-localizes together with GR at inflammatory promoters and enhancers in response to TLR4 signaling and glucocorticoids.

### GR recruits BRG1 to a distinct subset of macrophage binding sites

Since we had found protein-protein interactions and DNA co-occupancy between GR and the SWI/SNF complex, we performed ChIP-seq for the core component BRG1 in activated primary macrophages with and without GR ligand stimulation, to determine whether GR recruits BRG1 to chromatin. When analyzing the ∼8,000 GR binding sites shared with BRG1, we found that over 1,300 of them were dependent on GR ligand, meaning that BRG1 occupancy was induced by Dex in LPS-activated primary macrophages (**Fig. 2A**). Similar to previous studies for the GR co-regulators GRIP1 and SETD1A/COMPASS, we observed a ligand-mediated expansion of the BRG1 cistrome in macrophages (Uhlenhaut et al. 2013; Greulich et al. 2021b). Around 15,500 BRG1 binding sites were gained upon Dex stimulation, while ∼4,700 LPS-specific BRG1 sites were lost (Fig. S2A). Generally, most BRG1 binding sites were found in intronic or intergenic enhancer regions, under both conditions (Fig. S1B) (Hoffman et al. 2018).

**Figure 2:**
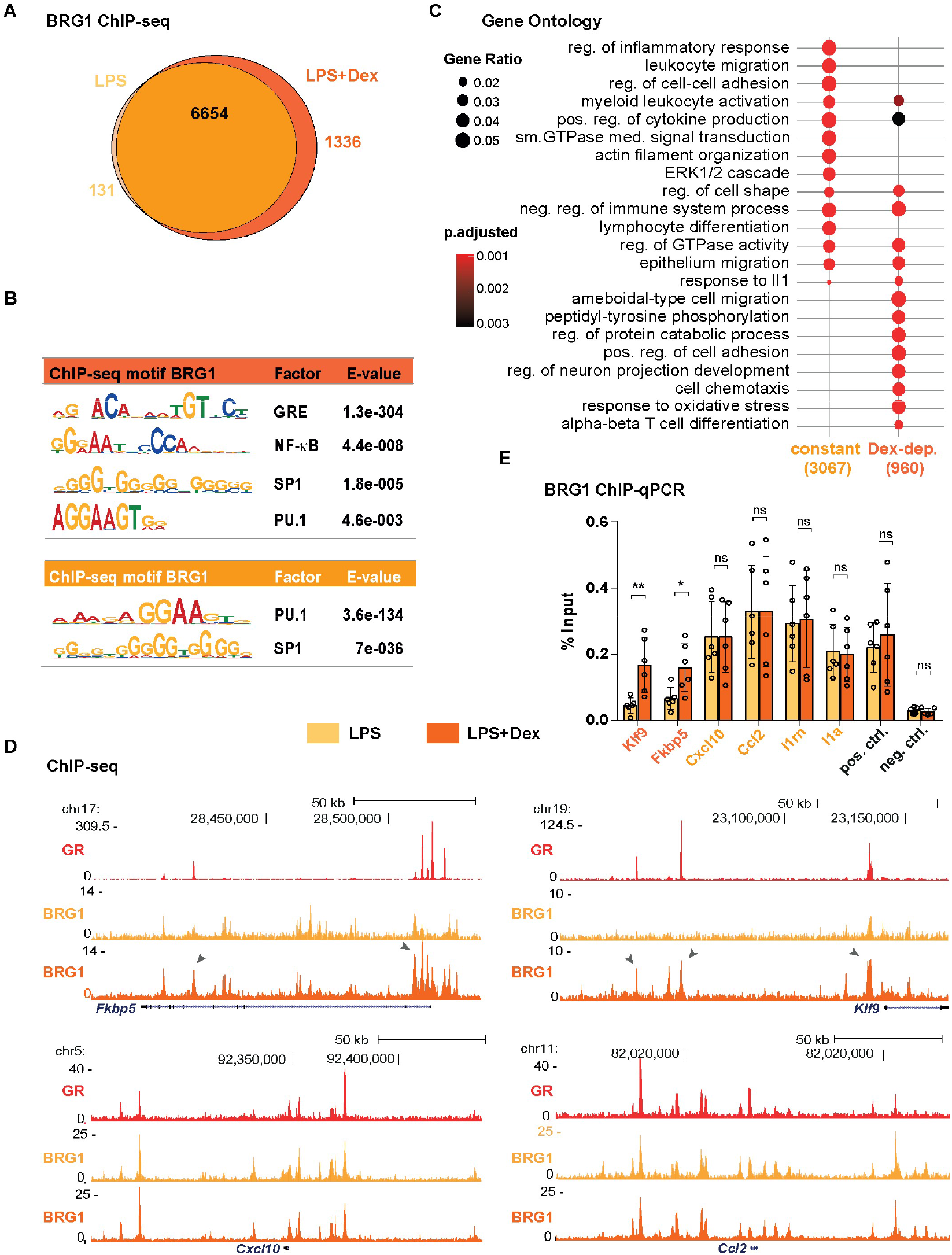
Locus-specific recruitment of BRG1 by GR in macrophages. (A) Venn diagram of BRG1 ChIP-seq peaks overlapping with the GR cistrome (∼8,000, see Fig. 1C), in response to LPS alone, or LPS plus Dexamethasone (Dex). (B) Motif enrichment of the 1,336 Dex-induced BRG1 peaks, and the constant BRG1 ChIP peaks (detected in both LPS and in LPS+Dex, 6,654) shown in A. (C) Functional annotation of the two BRG1 ChIP peak classes, Dex-induced and constant, based on the nearest gene. (D) Example genome browser tracks of GR and BRG1 ChIP-seq in macrophages treated with LPS alone (BRG1) or LPS plus Dex (GR, BRG1), means of n=2. Arrows point at sites of GR-induced BRG1 recruitment. (E) BRG1 ChIP-qPCR validation for selected loci shown in D. Error bars show standard deviation, * p<0.05, ** p<0.01, ns= not significant, unpaired two-tailed Student’s t-test, n=3.

Accordingly, these GR ligand-dependent BRG1 ChIP peaks featured a GRE consensus sequence as significantly enriched in motif analyses. Additional motifs include the ubiquitous, general transcription factor SP1, the master regulator of macrophage cell fate PU.1 and the inflammatory mediator NF-κB (**Fig. 2B**) (Glass and Natoli 2016). Of note, motifs for the inflammatory transcription factor AP-1 were identified in both BRG1 cistromes (LPS and LPS plus Dex), without a significant enrichment for the GR-BRG1 subset (Fig. S2C).

In line with GR’s prominent role in the transcriptional control of macrophage function and activity, these Dex-induced BRG1 binding sites mapped near genes involved in chemotaxis and migration, protein phosphorylation, metabolism and T cell activation (**Fig. 2C**) (Escoter-Torres et al. 2019). For example, both ChIP-seq as well as ChIP-qPCR show increased binding of BRG1 to the *Fkbp5* and the *Klf9 cis*-regulatory regions in response to Dex (**Fig. 2D&E**). These observations are consistent with transcriptional activation of *Fkbp5* and *Klf9*, for example, by GR, and with BRG1’s role in nucleosome remodeling and transcription by nuclear receptors (Trotter and Archer 2008).

Importantly, the majority of GR and BRG1 co-bound loci, which are associated with inflammatory pathways, were pre-bound by BRG1 in the absence of Dex, in line with their known function in LPS-activated macrophages (**Fig. 2C**, Fig. S2D). That means we did not detect changes in BRG1 ChIP-seq signal intensity between the samples treated with LPS only, and those treated with LPS plus Dex. For example, GR binding sites near *Ccl2, Cxcl10, Il1a* or *Il1rn* displayed robust BRG1 occupancy in both conditions (LPS and LPS+Dex) (**Fig. 2D&E**, Fig. S2E). Since these genes are expressed in LPS-activated macrophages, they may depend on the SWI/SNF complex for their induction upon TLR4 stimulation (McAndrew et al. 2016; Chen et al. 2020). Our observations suggest that GR does not appear to evict BRG1 in order to repress the transcription of chemokines, cytokines, interleukins *etc*., since we did not observe a significant reduction in global BRG1 occupancy in response to Dex, but rather a gain at specific activated GR target loci.

### Chromatin remodeling in response to GR ligand

As we had observed co-occupancy and recruitment of the core SWI/SNF subunit BRG1 at GR-bound *cis*-regulatory sites in murine macrophages, we performed ATAC-seq in LPS and in LPS+Dex treated cells, to measure chromatin accessibility in response to GR ligand. Overall, we identified over 100,000 sites of open chromatin in our primary macrophages, of which 8,860 were only present in macrophages treated with both LPS and Dex (**Fig. 3A**). Amongst those accessible regions, ∼27,800 displayed BRG1 occupancy and ∼8,200 showed GR co-binding, in LPS and Dex stimulated cells. Conversely, essentially all (99.8%) GR plus BRG1 co-occupied sites mapped to accessible chromatin (Fig. S3A).

**Figure 3:**
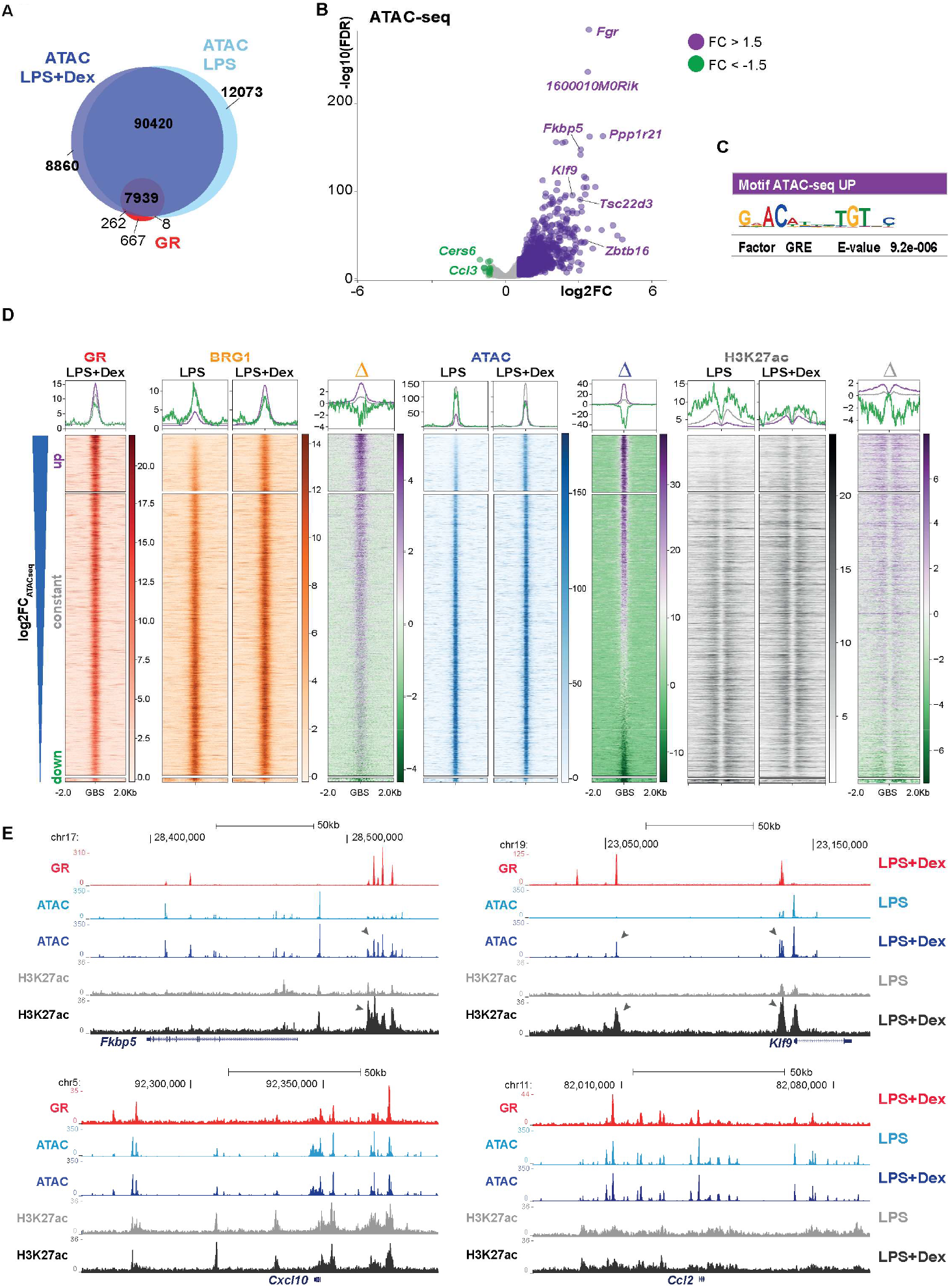
GR-induced macrophage chromatin accessibility changes. (A) Venn diagram with numbers of ATAC-seq peaks called in LPS and in LPS+Dex treated macrophages (n=4) and GR ChIP-seq (n=2). (B) Volcano plot of differential ATAC-seq signals at GR and BRG1 co-occupied regions, fold changes (FC) in LPS+Dex versus LPS treated samples. Dots represent single genomic regions, associated to the nearest gene. 7,351 peaks, n=4, FDR<0.05 (C) Differential motif enrichment analysis of the three categories of ATAC-seq peaks (shown in B), versus the union of all three peak sets (5,519 peaks). No significant enrichment was found for the gray or green sets. (D) Heatmaps of mean GR, BRG1, H3K27ac ChIP- and ATAC-seq signals at +/-2kb of GR-BRG1 co-occupied regions after LPS or LPS+Dex treatment. Sites are sorted by the Dex-induced change in ATAC-seq signal in descending order, and clustered by GR-BRG1 binding sites gaining (FC>1.5, FDR<0.05), maintaining (1.5>FC<-1.5, FDR>0.05) or losing (FC<-1.5, FDR<0.05) accessibility. Differential heatmaps (Δ) compare LPS+Dex versus LPS. Coverage plots on top summarize the median signal per group (GBS: GR binding site). (E) Representative genome browser tracks showing the mean signal of GR (n=2), ATAC-seq (n=4) and H3K27ac ChIP-seq (n=2) for *Fkbp5, Klf9, Ccl2* and *Cxcl10* loci. Arrows highlight signal changes.

When quantifying and comparing the ATAC-seq signal detected at *cis*-regulatory sites occupied by both GR and BRG1, we found that 1,234 loci gained ATAC-seq signals, while only 12 loci lost DNA accessibility. For example, classical GR target genes like *Fkbp5, Tsc22d3* (*Gilz*) and *Klf9* gained chromatin accessibility together with BRG1 recruitment upon Dexamethasone exposure (**Fig. 3B**). Consistent with retained BRG1 occupancy, we found only minimal reductions in ATAC-seq signals at GR target loci, on the other hand, indicating that GR does not generally close chromatin to repress transcription.

When performing a differential motif enrichment search among those sequences mapping to loci which gained openness in response to GR ligand, we found the GRE consensus motif over-represented among the ATAC-seq signatures (**Fig. 3C**). These results might point towards GR’s role in nucleosome positioning or phasing, possibly via BRG1 recruitment. Our data underscore the broad requirement and central role of the BRG1-containing SWI/SNF remodeling complex for transcriptional activation by GR (Hoffman et al. 2018). Furthermore, general motif enrichment analyses of our ATAC-seq signatures revealed consensus sites for the myeloid lineage factor PU.1 and the architectural factor CTCF, both of which are known to shape the macrophage chromatin landscape (Fig. S3B) (Ghirlando and Felsenfeld 2016).

**Fig. 3D** compares the BRG1 and the H3K27acetyl ChIP-seq reads with the ATAC-seq signal strength between LPS and LPS plus Dex treated macrophages, for all GR/BRG1 co-bound sites with either gained, reduced or constant (1.5>FC<-1.5, FDR>0.05) ATAC-seq signals (5,519 peaks in total). Generally, chromatin accessibility correlated with BRG1 recruitment and histone H3K27 acetylation induced by GR (**Fig. 3D**). Moreover, GR/BRG1 co-occupied loci with constant DNA accessibility were associated with genes involved in inflammation, such as ‘positive regulation of cytokine production’, ‘ERK1/2 cascade’ or ‘negative regulation of immune system processes’ (Fig. S3C).

For example, the *Klf9* and the *Fkbp5* loci both showed increased BRG1 occupancy, increased ATAC-seq read signals and increased histone H3K27 acetylation in response to Dex (**Fig. 3E**). Negative GR targets such as *Ccl2, Cxcl10, Il1a* or *Il1rn*, however, appeared to maintain a rather constant level of BRG1 binding, chromatin accessibility and H3K27 acetylation (**Fig. 3D**, Fig. S3D).

In general, our ATAC-seq profiling in primary macrophages revealed a cluster of distinct GR target loci, which displayed increased chromatin accessibility coinciding with ligand-activated BRG1 recruitment. Furthermore, a large fraction of GR-BRG1 co-bound genomic sites appeared to retain a constant level of BRG1 occupancy and openness not affected by GR ligand. The former subset mainly appears to correspond to activated GR target genes, while the latter seems to represent genes repressed by GR.

### BRG1 is required for transcriptional activation and repression by GR

Since our ChIP-seq and ATAC-seq profiles had exposed interactions between GR and the SWI/SNF complex at macrophage *cis*-regulatory elements, which manifested as BRG1 recruitment or co-occupancy together with chromatin remodeling or openness, respectively, we next performed loss of function studies. We knocked down *Brg1* expression in primary macrophages by siRNA, and performed RNA-seq to study the effects of *Brg1* inactivation on GR target gene regulation. Indeed, in macrophages treated with LPS and Dex, *Brg1* knockdown resulted in both up- and down-regulation of GR target genes compared to controls. For example, *Fkbp5, Klf9* and other positive GR targets were downregulated (induced to a lesser extent) upon transfection with *Brg1* siRNAs (**Fig. 4A**, Fig. S4A). Strikingly, many negative, inflammatory GR targets, such as *Ccl2, Ccl4, Cxcl10, Mmp27, Btg1, Il1a, Il1b* and *Il1rn* etc. were upregulated, meaning those were de-repressed. Generally, functional annotation of significantly differentially expressed genes showed an enrichment of genes involved in inflammation, immune responses, cytokines, defense responses and migration among those derepressed genes (**Fig. 4B**, Fig. S4B).

**Figure 4:**
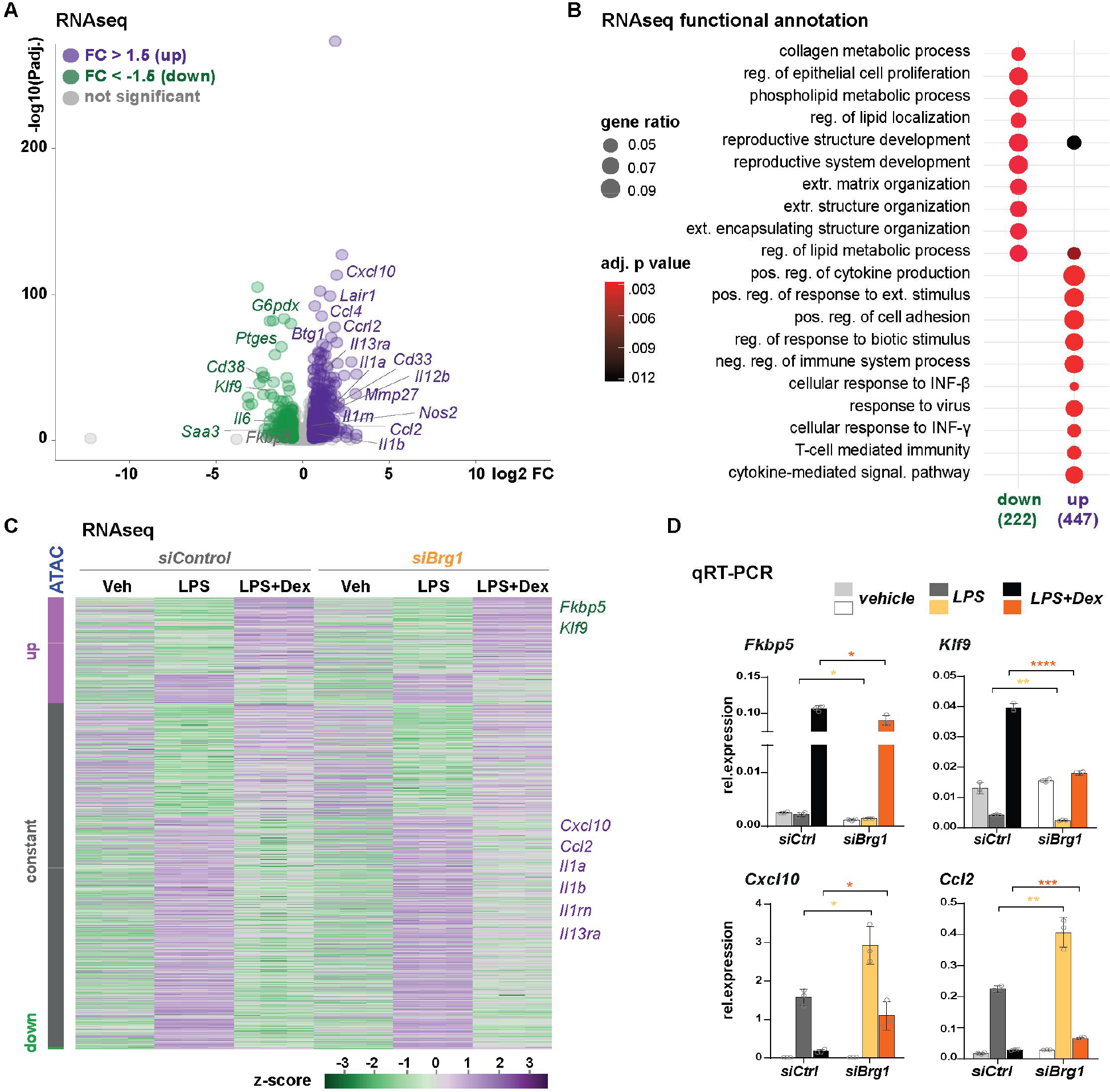
Loss of *Brg1* affects the glucocorticoid response in macrophages. (A) Volcano plot for transcripts harboring a nearby GR/BRG1 ChIP peak, showing RNAseq expression changes between control and *Brg1* knockdown macrophages treated with LPS+Dex (n=3, fold change ±1.5, p adjusted<0.05). (B) Gene Ontology enrichment (‘biological process’) of the differentially expressed common GR/BRG1 target genes shown in A. (C) Heatmap for GR/BRG1 targets associated with the three ATAC-seq categories (Fig. 3D), in control and *Brg1* knockdown macrophages treated with vehicle (Veh), LPS and LPS+Dex. (D) qRT-PCR validation of two positive and two negative GR/BRG1 targets upon *Brg1* or control siRNA transfection. Error bars show standard deviation, *p<0.05, **p<0.01, ***p<0.001, ****p<0.0001, ns = not significant, unpaired two-tailed Student’s t-test, n=3.

Importantly, with respect to the LPS response, many of these genes did not appear to depend on BRG1 for their activation by TLR4 signaling (**Fig. 4C**). Compared to quiescent macrophages, several inflammatory mediators were still induced upon LPS stimulation in *Brg1* knockdown samples. These effects were neither due to differential mRNA expression of the *GR* gene itself, nor downregulation of known GR co-regulators such as *GRIP1* or *Setd1a* (Fig. S4C&D).

Our RNA-seq profiles revealed that BRG1 is not only required for the transcriptional activation of nuclear receptor target genes, but also for the transcriptional repression of key inflammatory targets by GR. For example, *Cxcl10* and *Ccl2* were potently upregulated in *Brg1* knockdown and control cells activated by LPS, but showed impaired repression by Dex in the absence of BRG1 (**Fig. 4D**).

Taken together, our *Brg1* loss of function studies demonstrated a functional requirement of this enzymatic subunit not only for transcriptional activation, but also for transcriptional repression by GR, which could conceivably occur independently of its function in chromatin accessibility (**Fig. 4C**).

### BRG1 is required for histone deacetylation by GR

As we had observed impairments in both transcriptional activation and repression of GR target genes after *Brg1* siRNA knockdown in macrophages, we next aimed to validate these observations and to functionally characterize these affected loci. We first treated primary macrophages with a commercially available allosteric dual brahma homolog (BRM)/(BRG1) ATPase activity inhibitor (Papillon et al. 2018): As shown in **Fig. 5A**, inhibiting BRG1 catalytic activity reproducibly impaired the transcriptional activation of *Fkbp5* and *Klf9*, and compromised the transcriptional repression of *Ccl2, Cxcl10, Il1a* and *Il1rn* by GR in LPS-activated cells.

**Figure 5:**
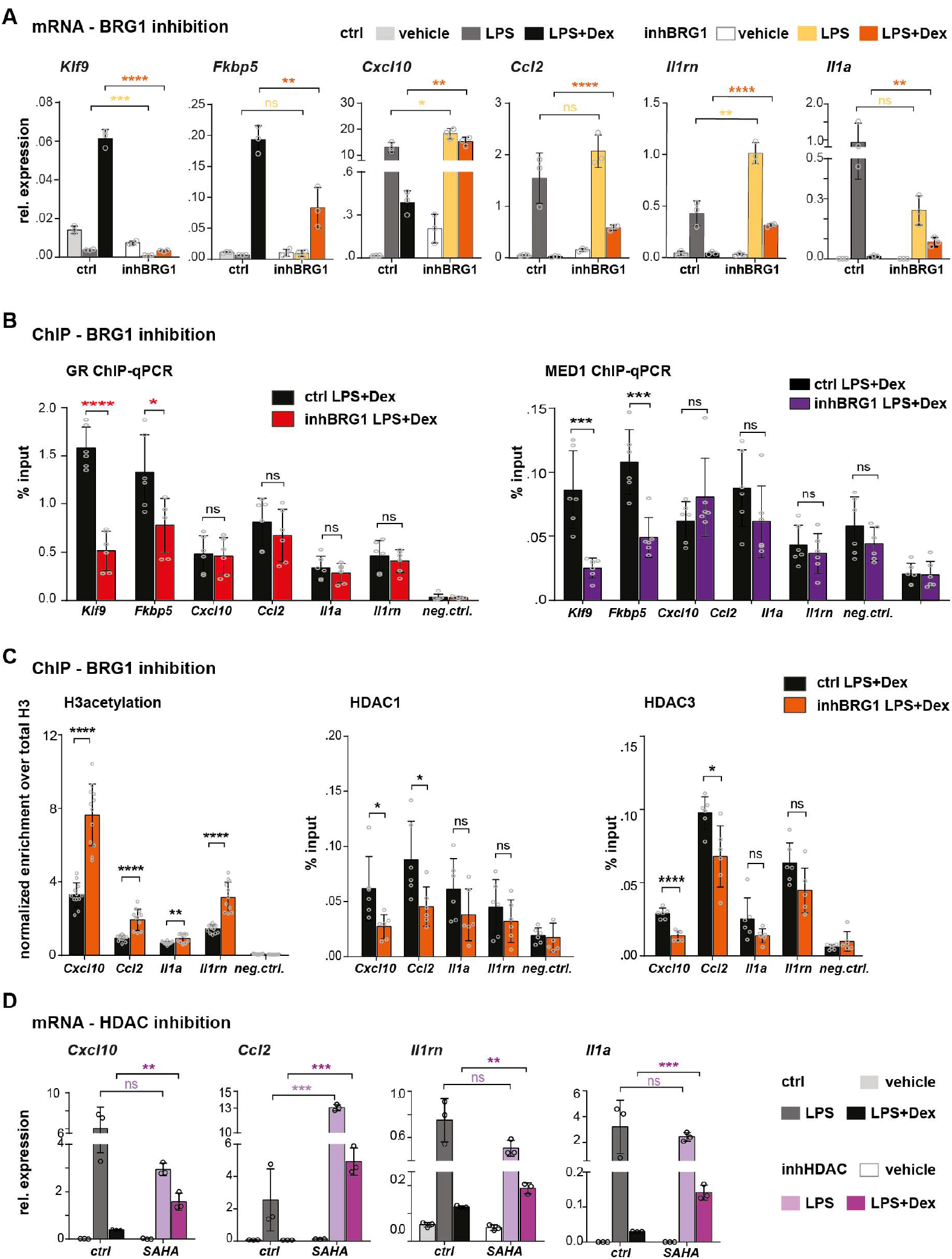
BRG1 catalytic activity is important for macrophage GR target gene regulation. (A) qRT-PCR of GR target genes in vehicle, LPS and LPS+Dex treated primary macrophages upon BRG1 inhibition (compared to DMSO). (B) GR and Mediator (Med1) ChIP-qPCR in control and BRG1 inhibitor treated macrophages (LPS+Dex). (C) Total histone H3 acetylation, HDAC1 and HDAC3 ChIP-qPCR in control and BRG1 inhibitor treated macrophages, upon LPS+Dex stimulation, at repressed GR target sites. For H3 acetylation ChIP, data were spike-in normalized and the values represent the % input over the total histone H3. (D) qRT-PCR of repressed GR target genes in vehicle, LPS and LPS+Dex stimulated macrophages treated with control (DMSO) or SAHA (HDAC inhibitor). For all bar graphs, values are mean ± standard deviation. *p<0.05, **p<0.01, ***p<0.001, ****p<0.0001, ns = not significant, unpaired two-tailed Student’s t-test, n=3 biological replicates.

When performing ChIP-qPCR for GR itself, in the presence of the SWI/SNF inhibitor, we found strongly reduced binding of the receptor to the *cis*-regulatory regions of the *Klf9* and *Fkbp5* genes, while the occupancy of the *Ccl2, Cxcl10, Il1a* and *Il1rn* binding sites was not affected (**Fig. 5B**). The diminished GR target gene binding and transcriptional activation was accompanied by weakened recruitment of the Mediator complex, as determined by ChIP-qPCR for the central MED1 subunit, at the *Fkbp5* and *Klf9* loci (Chen and Roeder 2007).

Conversely, *Cxcl10, Ccl2, Il1a* and *Il1rn*, which displayed impaired transcriptional repression by GR despite maintained chromatin interactions, showed increased total histone H3 acetylation correlating with increased mRNA production (**Fig. 5C**). These histone acetylation marks coincided with diminished recruitment of the histone deacetylases HDAC1 and HDAC3 in response to GR ligand. Of note, this observation refers to specific loci, as global HDAC activity was not diminished in primary macrophages treated with the BRG1 inhibitor (Fig. S5A).

To support our hypothesis that BRG1 might be required for the assembly of a functional corepressor complex containing HDACs and affecting the histone acetylation levels of inflammatory genes controlled by GR, we treated macrophages with the histone deacetylase inhibitor ‘Vorinostat’, also known as suberanilohydroxamic acid (SAHA) (Marks and Breslow 2007). Indeed, HDAC inhibition was able to recapitulate the impaired repression of *Ccl2, Cxcl10, Il1a* and *Il1rn* by GR, in macrophages treated with LPS and Dex (**Fig. 5D**). Importantly, these differential gene expression and chromatin pattern changes were observed despite maintained GR, BRG1, HDAC1, HDAC2 and HDAC3 mRNA and protein expression levels in these cells, and despite comparable BRG1 occupancy of these loci (Fig. S5B-F).

In conclusion, we found that BRG1 activity is essential for both transcriptional activation and repression of macrophage GR target genes. Our data may suggest that the transcriptional repression of inflammatory cytokines, chemokines and interleukins in response to glucocorticoids requires BRG1 for the assembly of a functional, HDAC-containing co-repressor complex. Conceivably, these findings point towards a novel role for the SWI/SNF complex independent of its nucleosome remodeling function.

## DISCUSSION

Our study revealed a dual role for the BRG1-containing SWI/SNF chromatin remodeling complex in GR-mediated inflammatory gene regulation in murine macrophages. Near activated GR target genes (such as *Klf9* and *Fkbp5*), we found that BRG1 was required for stable GR DNA binding and Mediator recruitment, coincident with increased chromatin accessibility. This continuous requirement of BRG1 for enhancer maintenance, openness and transcriptional activation is in line with previous reports on SWI/SNF complexes in other cell types (Hoffman et al. 2018; Iurlaro et al. 2021; Schick et al. 2021).

However, near negative GR target genes (like *Ccl2, Cxcl10, Il1rn and Il1a*), on the other hand, BRG1’s catalytic activity was necessary for transcriptional repression, independently of its chromatin remodeling function. For those loci, we found that the histone H3 acetylation levels were maintained after stimulation with Dexamethasone, rather than decreased, which concurred with increased mRNA expression (i.e., reduced repression). While the DNA accessibility remained constant, the impaired repression could be explained by reduced recruitment of the histone deacetylases HDAC1 and HDAC3 in response to GR ligand, especially since pharmacological HDAC inhibition mirrored this phenotype.

Interestingly, a requirement of BRG1 and HDAC2 for nuclear hormone receptor-mediated transcriptional repression was also shown for the closely related progesterone and estrogen receptors (Jung et al. 2001; Nacht et al. 2016). Furthermore, BRG1 was found to be critical for the formation of stable complexes between GR and HDAC2 on the *POMC* promoter, along with histone H4 de-acetylation and GR-dependent repression (Bilodeau et al., 2006).

SWI/SNF chromatin remodeling complexes have been described as having both co-activator as well as co-repressor functions and thus may provide a molecular hub or platform, switching from transcriptional activation to repression (Zhang et al. 2007; Kim et al. 2021). For example, locus-specific phosphorylation of BRG1 at Ser1382 has been reported to release the HDAC1/2-containing NURD complex and to favor BRG1’s nucleosome remodeling activity (Kim et al. 2021).

Currently, the molecular mechanisms that specify positive versus negative gene regulation by GR, mediated via co-activator or co-repressor complex assembly, respectively, remain elusive. Besides an enrichment for classical, palindromic GRE consensus motifs amongst GR binding sequences associated with *de novo* BRG1 recruitment and increased chromatin accessibility, we have not yet been able to identify discriminatory signatures or sequence motifs. It is conceivable that BRG1 represents a key interaction partner of GR, which might switch between co-activator and co-repressor conformations in a locus-specific manner, depending on the chromatin context.

In summary, our findings show that BRG1 is involved in anti-inflammatory glucocorticoid responses, which might suggest that future therapeutic approaches using SWI/SNF or HDAC inhibitors may have immunomodulatory effects.

## METHODS

### Cell lines

RAW264.7 cells (ATCC TIB-71, RRID CVCL 0493) were cultured in DMEM supplemented with 10% FBS and 1% penicillin/streptomycin. The cells were grown at 37 °C in the presence of 5% CO_2_.

*Drosophila* S2 cells (donated from P. Becker, RRID: CVCL_IZ06) were cultured in Schneider’s *Drosophila* medium supplemented with 10% FBS and 1% penicillin/streptomycin. The cells were grown in T175 flasks at 28 °C in absence of CO_2_.

### Extraction and differentiation of bone marrow derived macrophages

Leg bones were surgically removed from 6-14 weeks old wild type C57BL6/J male mice. After muscle dissection and clean-up of the bones with ethanol, bone marrow was extracted in RPMI. Erythrocytes were lysed with AKC lysis buffer (1 M NH_4_Cl, 1 M KHCO_3_, 0.5 M EDTA). Afterwards the cells were purified on a Ficoll-Paque gradient and cultured in differentiation medium (DMEM containing 30% supernatant of L929 cells, 20% FBS 1% penicillin/streptomycin) for 7 days on non - cell culture treated plates. Versene was applied to the differentiated macrophages, which were subsequently counted and seeded in macrophage serum free medium.

Cells were treated either with vehicle (0.1% EtOH and PBS), LPS (100 ng/ml, Sigma Aldrich and 0.1% EtOH) or LPS+Dex (100 ng/ml LPS Sigma; 1 µM Dexamethasone in EtOH). For the inhibitor experiments, macrophages were additionally treated either with 500 nM BRG1/BRM inhibitor (MedChemExpress, HY-119374) or with 1 µM SAHA (Sigma, SML0061) or 0.05%-0.1% DMSO, respectively, for 6 hours.

### Nuclear extraction and co-IP

RAW264.7 cells were treated with 1µM Dexamethasone overnight, followed by 3 hours treatment with 100ng/mL of LPS. The cells were washed thoroughly with ice-cold PBS and then lysed in V1 lysis buffer (10 mM HEPES-KOH pH 7.9, 1.5 mM MgCl_2_, 1 0mM KCl and freshly added 1 μM Dexamethasone, 0.5 mM DTT, 0.15% NP40, protease inhibitors and PhosphoSTOP) in a glass douncer on ice. After centrifugation at 2,700g for 20 min, the nuclei were collected and lysed in V2 buffer (420 mM NaCl, 20 mM HEPES-KOH pH 7.9, 20% glycerol, 2 mM MgCl_2_, 0.2 mM EDTA and freshly added 1 μM Dexamethasone, 0.5 mM DTT, 0.1% NP40, protease inhibitors and PhosphoSTOP) for 1 hour while agitating at 4°C. The nuclear extracts were collected after 45 min centrifugation at 21,000g at 4°C and used for co-IPs.

Co-IPs were performed with 200µg of nuclear protein extract that was pre-cleared with α-rabbit Dynabeads (Invitrogen) for 1 hour in IP buffer (20 mM Tris pH 8, 100 mM KCl, 5 mM MgCl_2_, 0.2 mM EDTA, 20% glycerol and freshly added protease inhibitors) under rotation at 4°C. The pre-cleared protein extracts were incubated with 3µg rabbit α-BRG1 (Cell Signalling, 49360), rabbit α-GR (Proteintech, 24050-1-AP), rabbit α-Baf57 (Bethyl Labs, A300-810A) and rabbit α-Baf60a (Proteintech, 10998-2-AP) antibody or 3µg of rabbit IgG antibody (Cell Signalling, 2729) for 2 hours under rotation at 4°C, followed by an overnight incubation with BSA blocked α-rabbit Dynabeads (Invitrogen) at 4°C. Beads were washed 3 times with IP buffer supplemented with 0.3% Triton X-100. Bound proteins were eluted in Laemmli buffer and DTT for 30 min at 37°C and analyzed by Western Blot using mouse α-GR (Santa Cruz, sc-393232), mouse α-Brg1 (Cell Signalling, E9O6E) and goat α-Baf60a (Santa Cruz, sc-82778) antibodies.

### siRNA mediated gene silencing

Gene silencing in primary macrophages was performed using the RNAimax kit (Invitrogen) in a 12-well plate according to the manufacturer’s instructions. Briefly, in each well, 50nM of siRNA diluted in 165µL serum free medium were mixed with 2µl of RNAimax in 165µl serum free medium. After 20 min of incubation at room temperature, 430.000 BMDMs were added to each well and incubated for 48 hours. Macrophages were treated either with vehicle, LPS or LPS+Dex for 6 hours before collection. We used non-targeted scramble control (D-001206-14) or si*Smarca4*/*Brg1* (M-041135-01-0005) (Dharmacon, siGenome, SMARTpool) siRNAs.

### RNA extraction, cDNA synthesis and qRT-PCR

Total RNA was extracted from macrophages using the RNeasy Mini Kit (Qiagen) and 500ng of mRNA were reverse transcribed using the QuantiTect reverse transcription kit (Qiagen) following the manufacturer’s instructions. qPCR was performed on Viia 6/7 Real time PCR system using SYBR Green master mix (Life Technologies). The primers used are listed in Supplementary table 1. The expression was normalized to the house keeping gene *Rplp0*.

### RNA-sequencing

RNA-seq was performed in BMDMs after siControl and siBRG1 knockdown. The RNA quality was determined on an Agilent 2100 Bioanalyzer with the RNA 6000 Nano kit, following manufacturer’s instructions. Library preparation and rRNA depletion were conducted using the TruSeq stranded mRNA Library Prep kit (Illumina) starting with 1µg of total RNA for each sample. The libraries were sequenced on the Illumina HiSeq4000 machine.

### ChIP-seq

40 million primary macrophages were used for each ChIP. The cells were treated with 100ng/ml LPS and with 1µM Dexamethasone or 0.1% EtOH for 3 hours and then fixed with 2mM disuccinimidyl glutarate (DSG) for 30 min at 4°C and 1% formaldehyde for 10 min at room temperature. The IP was performed using 8µg of rabbit α-GR (24050-1-AP, Proteintech) and 16µg of rabbit α-BRG1 (Cell Signalling, 49360 and Abcam ab110641, 8µg each) as previously described (Uhlenhaut et al. 2013). The DNA was quantified via Qubit, and the enrichment was validated by qPCR. Libraries were performed with the Kappa Hyperprep kit (Roche) according to the manufacturer’s instructions and sequenced on an Illumina NovaSeq6000 machine. The H3K27ac ChIP-seq dataset was previously published in *Greulich et al. 2021b*.

### ChIP-qPCR

For ChIP-qPCR, 2 million BMDMs were used. The cells were treated with DMSO or BRG1/BRM inhibitor and LPS or LPS+Dex for 3 hours. ChIP was performed as described previously (Uhlenhaut et al. 2013). 1µg of antibody was used for H3ac (Active Motif, 61937) and total H3 (Abcam, ab1791) IPs, and 2µg for BRG1 (Cell Signaling 49360 and Abcam ab110641, 1µg each), GR (24050-1-AP, Proteintech), MED1 (Bethyl labs, A300-793A), HDAC1 (Abcam, ab7028) and HDAC3 (Active Motif, ACM-40968) IPs. A spike-in normalization strategy with *Drosophila* chromatin was applied for the H3ac and total H3 IPs (Greulich et al. 2021a). qPCRs were performed with SYBR Green in a ViiA6/7 real time PCR system, and the enrichment was calculated as % input. H3ac samples were additionally normalized to total H3. The primers are listed in Supplementary table 2.

### ATAC-sequencing

For ATAC-seq, 50.000 BMDMs were treated either with 100ng/ml LPS or PBS, and 1µM Dexamethasone or 0.1% ethanol for 3 hours. Transposition was performed using the OmniATAC protocol (Corces et al. 2017) and the tagment DNA TDE1 enzyme (Illumina, 20034197). DNA was purified using the MinElute PCR purification kit (Qiagen). Afterwards, the transposed DNA was amplified using custom primers as previously described (Buenrostro et al. 2013). Libraries were purified using the MinElute PCR purification kit (Qiagen) and size selected for fragments 150bp-600bp using the Agencourt AMPure XP beads (Beckman Coulter). The quality of the libraries was determined by the Qubit dsDNA HS kit (Thermo Scientific) and the Agilent High Sensitivity DNA 2100 Bioanalyzer. The samples were sequenced on an Illumina Novaseq 6000 machine.

### Western blot

Nuclear extraction was performed in LPS+Dex primary macrophages treated either with DMSO control or BRG1 inhibitor or SAHA as described above. Western blot was performed using standard procedures with the following antibodies: mouse α-BRG1 (Cell Signalling, 52251), rabbit α-GR (Cell Signalling,12041), rabbit α-SNRP70 (Abcam, ab83306), mouse α-HDAC1 (Cell Signalling, 5356), mouse α-HDAC2 (Cell signalling, 5113) and mouse α-HDAC3 (Cell Signalling, 3949).

### HDAC activity assay

HDAC activity assays were performed using the HDAC GLO I/II assay kit (Promega, G6430) in 96 well plates following the manufacturer’s instructions. BMDMs were seeded in phenol-red free DMEM (Gibco, 21063-029), stimulated with LPS plus Dex and treated either with DMSO or with BRG1 inhibitors or SAHA as described above.

### NGS data analysis

NGS data quality was assessed with FastQC (RRID:SCR 014583, http://www.bioinformatics.babraham.ac.uk/projects/fastqc/).

For RNA Sequencing, gene-level quantification was performed with Salmon version 1.4.0 (RRID:SCR_017036 (Patro et al. 2017)). Settings were: -libType A, -gcBias, -biasSpeedSamp 5 using the mm10 (M25, GRCm38, mm10) reference transcriptome provided by Genecode (Frankish et al. 2019). Gene count normalization and differential expression analysis was performed with DESeq2 version 1.32.0 (RRID:SCR_015687 (Love et al. 2014)) after import of gene-level estimates with “tximport” version 1.20.0 (RRID:SCR_016752 (Soneson et al. 2015)) in R (RRID:SCR_001905, R version 4.1.0 (Team 2017)).

For gene annotation, Ensembl gene Ids were mapped to MGI symbols using the Bioconductor package “biomaRt” version 2.48.2 (RRID:SCR_002987 (Durinck et al. 2009)) and genome information was provided by Ensembl (GRCm38.p6 (Cunningham et al. 2019)). Genes with at least 1 read count, fold change of 1.5 and Benjamini-Hochberg-adjusted p-value < 0.05 were called significantly changed. We compared BMDMs after *Brg1* and control siRNA knockdown under LPS+Dex conditions (Table S3). Plots were generated with “ggplot2” version 3.3.5 (RRID:SCR_014601, (Wickham 2016)) or “pheatmap” version 1.0.12 (RRID:SCR_016418, https://github.com/raivokolde/pheatmap) packages and GO enrichment performed with “clusterProfiler” version 3.18 (RRID:SCR 016884 (Yu et al. 2012)) (Table S4). Details on the downstream analysis is documented in the R scripts available on github (https://github.com/FranziG/GRandBrg1).

ChIP-seq and ATAC-seq paired-end reads were mapped to the murine reference genome mm10 (Ensembl GRCm38.p6 (Cunningham et al. 2019)) with BWA-MEM version 0.7.13 (RRID:SCR 010910 (Li 2013)) or Bowtie2 version 2.4.2 (RRID: SCR 005476 (Langmead and Salzberg 2012)) respectively, and PCR duplicates were removed using Picard Tools version 2.0.1 (RRID:SCR -006525, http://picard.sourceforge.net/). Samples with duplication levels above 25% (ATAC-seq) or 50% (ChIP-seq) were excluded from further analysis. For visualization, bam files were filtered for properly paired and mapped reads and multimappers were removed with Samtools version 1.11 (RRID:SCR 002105 (Li et al. 2009)). Alignments were converted to bigwig files, merging 10 bp per bin using ‘bamCoverage’ from the Deeptools package version 3.5.0 (RRID:SCR -016366 (Ramirez et al. 2016)). Tracks were visualized with UCSC genome browser (Kent et al. 2002). Peaks were called with MACS version 3.0.0a5 in BAMPE mode and an FDR cutoff of 0.05. ChIP-seq peaks were called over matched input controls. Blacklisted regions (http://mitra.stanford.edu/kundaje/akundaje/release/blacklists/mm10-mouse/mm10.blacklist.bed.gz) were removed from analyses. Peak annotation was performed in R version 4.0.3 (RRID:SCR 014601 (Team 2017)) using the ChIPpeakAnno package version 3.24.1 and annotation data from the mouse Ensembl genome GRCm38.p6 (mm10 (Cunningham et al. 2019)).

The peak union of all replicates was used to determine reads in peaks (RiP) ratios and scaling factors to normalize for library size and background-to-noise ratio. Genome browser tracks were normalized by the RiP fraction.

For peak overlaps, reproducible peaks (peak intersection in at least 2 replicates) were used and displayed as Venn diagrams, made in R version 4.0.3 (RRID:SCR 014601 (Team 2017)) using the VennDiagram package version 1.6.20. Peaks regions were defined as overlapping when overlapping by at least 1bp using the GenomicRanges package version 1.42.0 (RRID:SCR 000025 (Lawrence et al. 2013)) in R. Peaks were annotated to the closest gene expressed in macrophages in any of our conditions with the ‘ChIPpeakAnno’ package version 3.24.1 (RRID:SCR 012828 (Zhu et al. 2010)) (Table S5). Genes were called expressed when passing a mean expression value of the 25th percentile. Enrichment analysis for Gene Ontology (GO) of Biological Processes was performed using the ‘clusterProfiler’ package 3.18.0 (RRID:SCR 016884 (Yu et al. 2012)) (Table S4). GO terms with more than 60% similarity in gene composition were removed, and only the term with the lowest Benjamini-Hochberg adjusted p-value was reported. Results of GO enrichment analyses are displayed as dot plots showing the top 20 enriched GO terms (by Benjamini-Hochberg adjusted p-value), sorted by gene ratio (proportion of set genes enriched in GO term). Motif enrichment was performed on peaks trimmed to 100 bp or 300 bp around the peak center with MEME suite version 5.3.0 (RRID:SCR 001783 (Machanick and Bailey 2011)) in enrichment or differential mode. MEME parameters were set to: ‘-dna –mod zoops -minw 5 -maxw 25 -nmotifs 20 -p 10’ using the JASPAR (2018 version, RRID:SCR - 003030 (Khan et al. 2018)), Uniprobe (RRID:SCR 005803 (Newburger and Bulyk 2009)) and SwissRegulon (RRID:SCR 005333 (Pachkov et al. 2013)) databases.

### Data access

Scripts and analytical details are available on github (https://github.com/FranziG/GRandBrg1). Previously published data for H3K27ac ChIP-seq in murine macrophages is accessible on GEO with the accession numbers GSM4040445-48.

All next generation sequencing data generated in this study is available on the NCBI Gene Expression Omnibus as a SuperSeries with the accession number GSE186514 (ATAC-seq: GSE186511, ChIP-seq: GSE1865112, RNA-seq: GSE1865113), Reviewer token: avmrmccqfjqnvyh.

## ACKNOWLEDGMENTS

We would sincerely like to thank T. Horn, S. Regn, I. Guderian, O. Garcia-Gonzalez and A. P. Syed for their contributions to this study. We are grateful for the help of I. de la Rosa Velazquez, B. Haderlein, the HMGU genomics core and the animal facilities (AVM). This project received funding from the Deutsche Forschungsgemeinschaft DFG (SFB 1064 Chromatin Dynamics Project ID 213249687 to NHU and to GS, TRR 205 Adrenal Research to NHU, Entzuendungsprozesse GR 5179/1-1 to FG and SFB 1321 Pancreatic Cancer Project ID 329628492 to GS) and from the ERC (ERC-2014-StG 638573 SILENCE to NHU).

## AUTHOR CONTRIBUTIONS

AM designed and performed the majority of the experiments together with NHU and FG. AM and CJ performed NGS experiments in primary macrophages, FG performed bioinformatics analyses, and BAS performed additional experiments. FMC and GS supported the establishment of ATAC-seq protocols. NHU secured funding, supervised the work, and wrote the manuscript together with AM and FG. All the authors participated in writing, reviewing and editing the manuscript.

## REFERENCES

Biddie SC, John S, Sabo PJ, Thurman RE, Johnson TA, Schiltz RL, Miranda TB, Sung MH, Trump S, Lightman SL et al. 2011. Transcription factor AP1 potentiates chromatin accessibility and glucocorticoid receptor binding. Mol Cell 43: 145–155.

Bilodeau S, Vallette-Kasic S, Gauthier Y, Figarella-Branger D, Brue T, Berthelet F, Lacroix A, Batista D, Stratakis C, Hanson J et al. 2006. Role of Brg1 and HDAC2 in GR trans-repression of the pituitary POMC gene and misexpression in Cushing disease. Genes Dev 20: 2871–2886.

Buenrostro JD, Giresi PG, Zaba LC, Chang HY, Greenleaf WJ. 2013. Transposition of native chromatin for fast and sensitive epigenomic profiling of open chromatin, DNA-binding proteins and nucleosome position. Nat Methods 10: 1213–1218.

Chen S, Yang J, Wei Y, Wei X. 2020. Epigenetic regulation of macrophages: from homeostasis maintenance to host defense. Cell Mol Immunol 17: 36–49.

Chen W, Roeder RG. 2007. The Mediator subunit MED1/TRAP220 is required for optimal glucocorticoid receptor-mediated transcription activation. Nucleic Acids Res 35: 6161–6169.

Corces MR, Trevino AE, Hamilton EG, Greenside PG, Sinnott-Armstrong NA, Vesuna S, Satpathy AT, Rubin AJ, Montine KS, Wu B et al. 2017. An improved ATAC-seq protocol reduces background and enables interrogation of frozen tissues. Nat Methods 14: 959–962.

Cunningham F, Achuthan P, Akanni W, Allen J, Amode MR, Armean IM, Bennett R, Bhai J, Billis K, Boddu S et al. 2019. Ensembl 2019. Nucleic Acids Res 47: D745–D751.

Durinck S, Spellman PT, Birney E, Huber W. 2009. Mapping identifiers for the integration of genomic datasets with the R/Bioconductor package biomaRt. Nat Protoc 4: 1184–1191.

Escoter-Torres L, Caratti G, Mechtidou A, Tuckermann J, Uhlenhaut NH, Vettorazzi S. 2019. Fighting the Fire: Mechanisms of Inflammatory Gene Regulation by the Glucocorticoid Receptor. Front Immunol 10: 1859.

Escoter-Torres L, Greulich F, Quagliarini F, Wierer M, Uhlenhaut NH. 2020. Anti-inflammatory functions of the glucocorticoid receptor require DNA binding. Nucleic Acids Res 48: 8393–8407.

Frankish A, Diekhans M, Ferreira AM, Johnson R, Jungreis I, Loveland J, Mudge JM, Sisu C, Wright J, Armstrong J et al. 2019. GENCODE reference annotation for the human and mouse genomes. Nucleic Acids Res 47: D766–D773.

Fryer CJ, Archer TK. 1998. Chromatin remodelling by the glucocorticoid receptor requires the BRG1 complex. Nature 393: 88–91.

Ghirlando R, Felsenfeld G. 2016. CTCF: making the right connections. Genes Dev 30: 881–891.

Glass CK, Natoli G. 2016. Molecular control of activation and priming in macrophages. Nature immunology 17: 26–33.

Greulich F, Hemmer MC, Rollins DA, Rogatsky I, Uhlenhaut NH. 2016. There goes the neighborhood: Assembly of transcriptional complexes during the regulation of metabolism and inflammation by the glucocorticoid receptor. Steroids 114: 7–15.

Greulich F, Mechtidou A, Horn T, Uhlenhaut NH. 2021a. Protocol for using heterologous spike-ins to normalize for technical variation in chromatin immunoprecipitation. STAR Protoc 2: 100609.

Greulich F, Wierer M, Mechtidou A, Gonzalez-Garcia O, Uhlenhaut NH. 2021b. The glucocorticoid receptor recruits the COMPASS complex to regulate inflammatory transcription at macrophage enhancers. Cell Rep 34: 108742.

Group RC, Horby P, Lim WS, Emberson JR, Mafham M, Bell JL, Linsell L, Staplin N, Brightling C, Ustianowski A et al. 2021. Dexamethasone in Hospitalized Patients with Covid-19. N Engl J Med 384: 693–704.

Hoffman JA, Trotter KW, Ward JM, Archer TK. 2018. BRG1 governs glucocorticoid receptor interactions with chromatin and pioneer factors across the genome. Elife 7.

Hsiao PW, Fryer CJ, Trotter KW, Wang W, Archer TK. 2003. BAF60a mediates critical interactions between nuclear receptors and the BRG1 chromatin-remodeling complex for transactivation. Mol Cell Biol 23: 6210–6220.

Iurlaro M, Stadler MB, Masoni F, Jagani Z, Galli GG, Schubeler D. 2021. Mammalian SWI/SNF continuously restores local accessibility to chromatin. Nat Genet 53: 279–287.

John S, Sabo PJ, Johnson TA, Sung MH, Biddie SC, Lightman SL, Voss TC, Davis SR, Meltzer PS, Stamatoyannopoulos JA et al. 2008. Interaction of the glucocorticoid receptor with the chromatin landscape. Mol Cell 29: 611–624.

John S, Sabo PJ, Thurman RE, Sung MH, Biddie SC, Johnson TA, Hager GL, Stamatoyannopoulos JA. 2011. Chromatin accessibility pre-determines glucocorticoid receptor binding patterns. Nat Genet 43: 264–268.

Jung DJ, Lee SK, Lee JW. 2001. Agonist-dependent repression mediated by mutant estrogen receptor alpha that lacks the activation function 2 core domain. J Biol Chem 276: 37280–37283.

Kent WJ, Sugnet CW, Furey TS, Roskin KM, Pringle TH, Zahler AM, Haussler D. 2002. The human genome browser at UCSC. Genome Res 12: 996–1006.

Khan A, Fornes O, Stigliani A, Gheorghe M, Castro-Mondragon JA, van der Lee R, Bessy A, Cheneby J, Kulkarni SR, Tan G et al. 2018. JASPAR 2018: update of the open-access database of transcription factor binding profiles and its web framework. Nucleic Acids Res 46: D1284.

Kim B, Luo Y, Zhan X, Zhang Z, Shi X, Yi J, Xuan Z, Wu J. 2021. Neuronal activity-induced BRG1 phosphorylation regulates enhancer activation. Cell Rep 36: 109357.

Langmead B, Salzberg SL. 2012. Fast gapped-read alignment with Bowtie 2. Nat Methods 9: 357–359.

Lawrence M, Huber W, Pages H, Aboyoun P, Carlson M, Gentleman R, Morgan MT, Carey VJ. 2013. Software for computing and annotating genomic ranges. PLoS Comput Biol 9: e1003118.

Li H. 2013. Aligning sequence reads, clone sequences and assembly contigs with BWA-MEM. arXiv:13033997.

Li H, Handsaker B, Wysoker A, Fennell T, Ruan J, Homer N, Marth G, Abecasis G, Durbin R, Genome Project Data Processing S. 2009. The Sequence Alignment/Map format and SAMtools. Bioinformatics 25: 2078–2079.

Love MI, Huber W, Anders S. 2014. Moderated estimation of fold change and dispersion for RNA-seq data with DESeq2. Genome Biol 15: 550.

Machanick P, Bailey TL. 2011. MEME-ChIP: motif analysis of large DNA datasets. Bioinformatics 27: 1696–1697.

Marks PA, Breslow R. 2007. Dimethyl sulfoxide to vorinostat: development of this histone deacetylase inhibitor as an anticancer drug. Nat Biotechnol 25: 84–90.

McAndrew MJ, Gjidoda A, Tagore M, Miksanek T, Floer M. 2016. Chromatin Remodeler Recruitment during Macrophage Differentiation Facilitates Transcription Factor Binding to Enhancers in Mature Cells. J Biol Chem 291: 18058–18071.

Muratcioglu S, Presman DM, Pooley JR, Grontved L, Hager GL, Nussinov R, Keskin O, Gursoy A. 2015. Structural Modeling of GR Interactions with the SWI/SNF Chromatin Remodeling Complex and C/EBP. Biophys J 109: 1227–1239.

Nacht AS, Pohl A, Zaurin R, Soronellas D, Quilez J, Sharma P, Wright RH, Beato M, Vicent GP. 2016. Hormone-induced repression of genes requires BRG1-mediated H1.2 deposition at target promoters. EMBO J 35: 1822–1843.

Newburger DE, Bulyk ML. 2009. UniPROBE: an online database of protein binding microarray data on protein-DNA interactions. Nucleic Acids Res 37: D77–82.

Pachkov M, Balwierz PJ, Arnold P, Ozonov E, van Nimwegen E. 2013. SwissRegulon, a database of genome-wide annotations of regulatory sites: recent updates. Nucleic Acids Res 41: D214–220.

Papillon JPN, Nakajima K, Adair CD, Hempel J, Jouk AO, Karki RG, Mathieu S, Mobitz H, Ntaganda R, Smith T et al. 2018. Discovery of Orally Active Inhibitors of Brahma Homolog (BRM)/SMARCA2 ATPase Activity for the Treatment of Brahma Related Gene 1 (BRG1)/SMARCA4-Mutant Cancers. J Med Chem 61: 10155–10172.

Patro R, Duggal G, Love MI, Irizarry RA, Kingsford C. 2017. Salmon provides fast and bias-aware quantification of transcript expression. Nat Methods 14: 417–419.

Quagliarini F, Mir AA, Balazs K, Wierer M, Dyar KA, Jouffe C, Makris K, Hawe J, Heinig M, Filipp FV et al. 2019. Cistromic Reprogramming of the Diurnal Glucocorticoid Hormone Response by High-Fat Diet. Mol Cell 76: 531–545 e535.

Ramirez F, Ryan DP, Gruning B, Bhardwaj V, Kilpert F, Richter AS, Heyne S, Dundar F, Manke T. 2016. deepTools2: a next generation web server for deep-sequencing data analysis. Nucleic Acids Res 44: W160–165.

Schick S, Grosche S, Kohl KE, Drpic D, Jaeger MG, Marella NC, Imrichova H, Lin JG, Hofstatter G, Schuster M et al. 2021. Acute BAF perturbation causes immediate changes in chromatin accessibility. Nat Genet 53: 269–278.

Soneson C, Love MI, Robinson MD. 2015. Differential analyses for RNA-seq: transcript-level estimates improve gene-level inferences. F1000Res 4: 1521.

Team RC. 2017. R: A Language and Environment for Statistical Computing. R Foundation for Statistical Computing, Vienna, Austria.

Trotter KW, Archer TK. 2004. Reconstitution of glucocorticoid receptor-dependent transcription in vivo. Mol Cell Biol 24: 3347–3358.

Trotter KW, Archer TK. 2008. The BRG1 transcriptional coregulator. Nucl Recept Signal 6: e004.

Trotter KW, King HA, Archer TK. 2015. Glucocorticoid Receptor Transcriptional Activation via the BRG1-Dependent Recruitment of TOP2beta and Ku70/86. Mol Cell Biol 35: 2799–2817.

Uhlenhaut NH, Barish GD, Yu RT, Downes M, Karunasiri M, Liddle C, Schwalie P, Hubner N, Evans RM. 2013. Insights into negative regulation by the glucocorticoid receptor from genome-wide profiling of inflammatory cistromes. Mol Cell 49: 158–171.

Wickham H. 2016. Elegant Graphics for Data Analysis. Springer-Verlag New York.

Yu G, Wang LG, Han Y, He QY. 2012. clusterProfiler: an R package for comparing biological themes among gene clusters. OMICS 16: 284–287.

Zhang B, Chambers KJ, Faller DV, Wang S. 2007. Reprogramming of the SWI/SNF complex for co-activation or co-repression in prohibitin-mediated estrogen receptor regulation. Oncogene 26: 7153–7157.

Zhu LJ, Gazin C, Lawson ND, Pages H, Lin SM, Lapointe DS, Green MR. 2010. ChIPpeakAnno: a Bioconductor package to annotate ChIP-seq and ChIP-chip data. BMC Bioinformatics 11: 237.

