## Supplemental Figures S1-S5 for "BRG1 defines a genomic subset of inflammatory genes transcriptionally controlled by the glucocorticoid receptor"

Mechtidou A. *et al.,*

**Supplementary Figures S1 - S5**

Supplementary Figure 1. GR interacts with SWI/SNF complex
Supplementary Figure 2. BRG1 ChIP-seq in macrophages (LPS and LPS+Dex)
Supplementary Figure 3. ATAC-seq in primary macrophages
Supplementary Figure 4. Loss of BRG1 affects GR function in macrophages
Supplementary Figure 5. Macrophage BRG1 and HDAC inhibition

**Supplementary Tables S1 - S5**

Supplementary Table S1: qRT-PCR primers
Supplementary Table S2: ChIP-qPCR primers
Supplementary Table S3: Differentially expressed genes (BMDMs, *siBRG1*, LPS+Dex)
Supplementary Table S4: Gene ontology annotation of various gene sets
Supplementary Table S5: Macrophage GR & BRG1 ChIP-seq plus ATAC-seq peaks

### Supplementary Figure 1

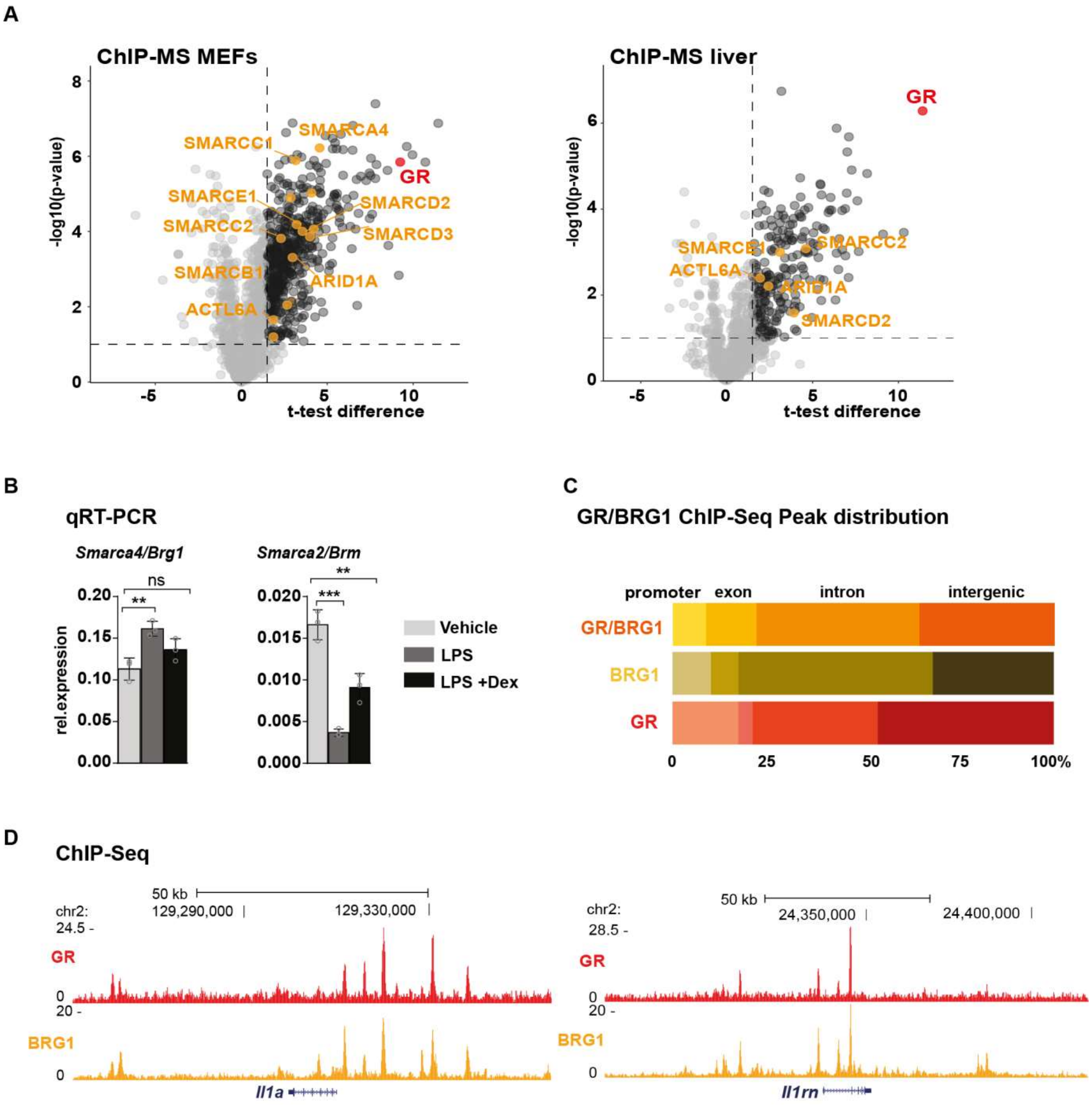


**Supplementary Figure 1. GR interacts with SWI/SNF complex.** (A) GR ChIP-MS interactomes including subunits of the SWI/SNF chromatin remodeling complex from mouse embryonic fibroblasts (MEFs stimulated with LPS and Dex) and from mouse livers. (Quagliarini et al. 2019; Escoter-Torres et al. 2020) SWI/SNF components are marked in orange. (B) qRT-PCR of *Smarca4/Brg1* and *Smarca2/Brm* in vehicle, LPS and LPS plus Dex stimulated macrophages. Bars = mean ± standard deviation, ns = not significant, unpaired two-tailed Student's t-test, n=3. (C) Genomic feature distribution of GR and BRG1-bound sites and regions specifically occupied by either BRG1 and/or GR. ChIP-seq peak sets as in Fig. 1C. (D) Example genome browser tracks for *Il1a* and *Il1rn* loci showing the mean signal from two GR and BRG1 ChIP-seq replicates in LPS plus Dex treated macrophages (n=2).

### Supplementary Figure 2

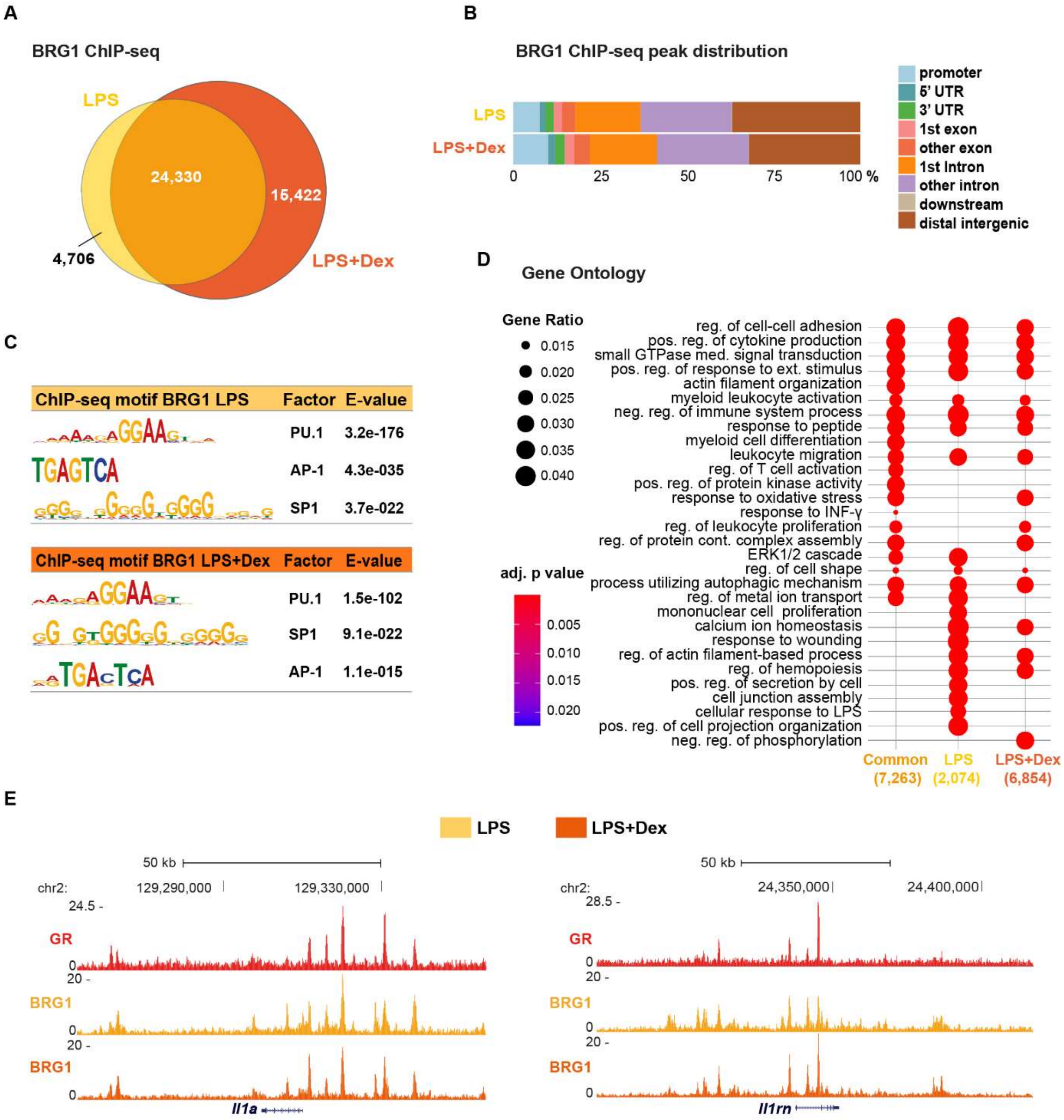


**Supplementary Figure 2. BRG1 ChIP-seq in macrophages treated with LPS and with LPS plus Dex.** (A) Overlap of BRG1 binding sites in macrophages treated with LPS plus Dex (n=2) or with LPS only (n=2). (B) Genomic feature distribution of BRG1-bound sites for both conditions. (C) MEME motif enrichment of BRG1 ChIP-seq peaks. Motifs were filtered for E<0.01. (D) Enrichment for biological processes of genes associated with nearby BRG1 binding sites, for common (orange) or unique (either LPS or LPS plus Dex) macrophage peaks. neg. = negative, pos. = positive, reg. = regulation (E) Representative genome browser tracks for *Il1a* and *Il1rn* loci showing the mean signal of GR and BRG1 ChIP-seq in macrophages (n=2 each).

### Supplementary Figure 3

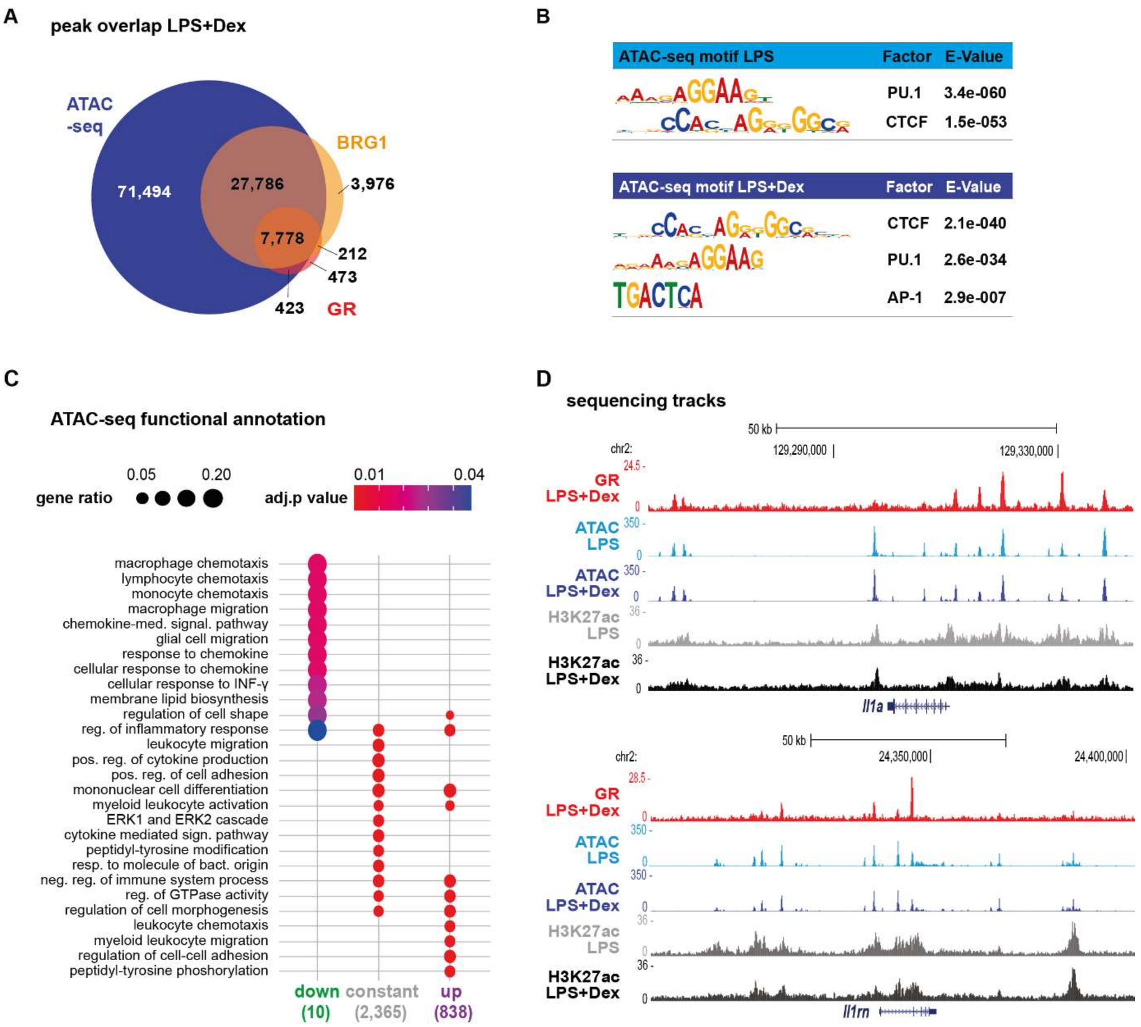


**Supplementary Figure 3. ATAC-seq in macrophages responding to LPS or LPS plus Dex.** (A) Overlap of accessible regions (ATAC-seq, n=4) with the BRG1 (n=2) and GR (n=2) binding sites, as determined by ChIP-seq in LPS+Dex treated macrophages. (B) MEME motif enrichment for the macrophage ATAC-seq peaks. Motifs were filtered for E<0.01. (C) Gene Ontology enrichment (biological process) of genes associated with the three categories of ATAC-seq peaks shown in Fig. 3B. cell. = cellular, neg. = negative, pos. = positive, reg. = regulation, resp. = response. (D) Representative genome browser tracks for *Il1a* and *Il1rn* loci showing mean ATAC-seq (n=4), GR (n=2) and H3K27ac ChIP-seq (n=2) coverage.

### Supplementary Figure 4

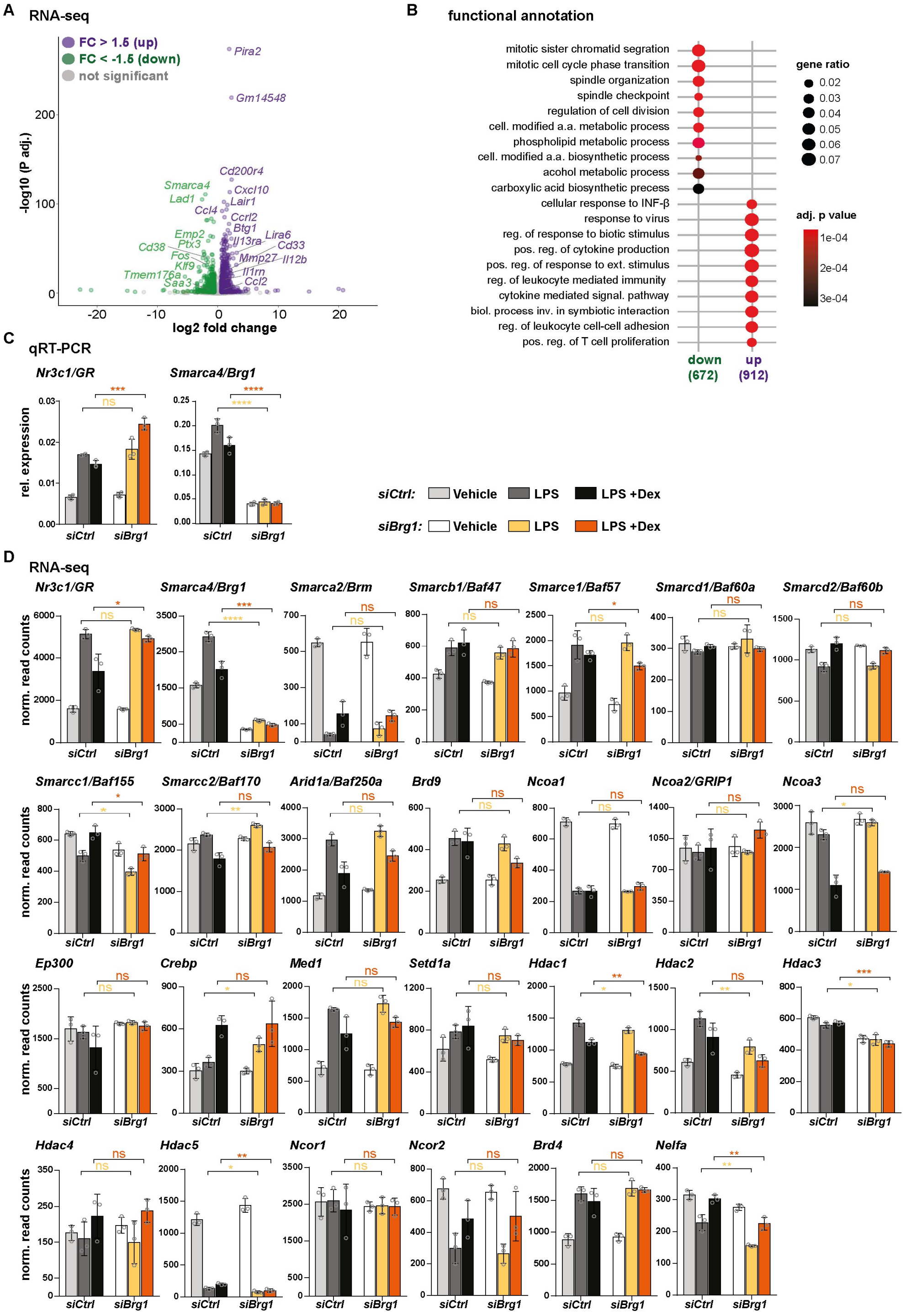


**Supplementary Figure 4. *Brg1* knockdown affects macrophage gene expression.** (A) Transcripts with differential expression between control and *Brg1* knockdown macrophages upon LPS+Dex treatment. (n=3, fold change ±1.5, p adjusted <0.05). (B) Gene Ontology enrichment for 'biological process' of differentially regulated genes in A. (C) qRT-PCR for *GR* and *Brg1* in control and *Brg1* knockdown macrophages. (D) DESeq-normalized RNAseq read counts for relevant factors in control and *Brg1* knockdown macrophages. For all bar graphs, values are means ± standard deviation. *p<0.05, **p<0.01, ***p<0.001, ****p<0.0001, ns= not significant, unpaired two-tailed Student's t-test, n=3.

### Supplementary Figure 5

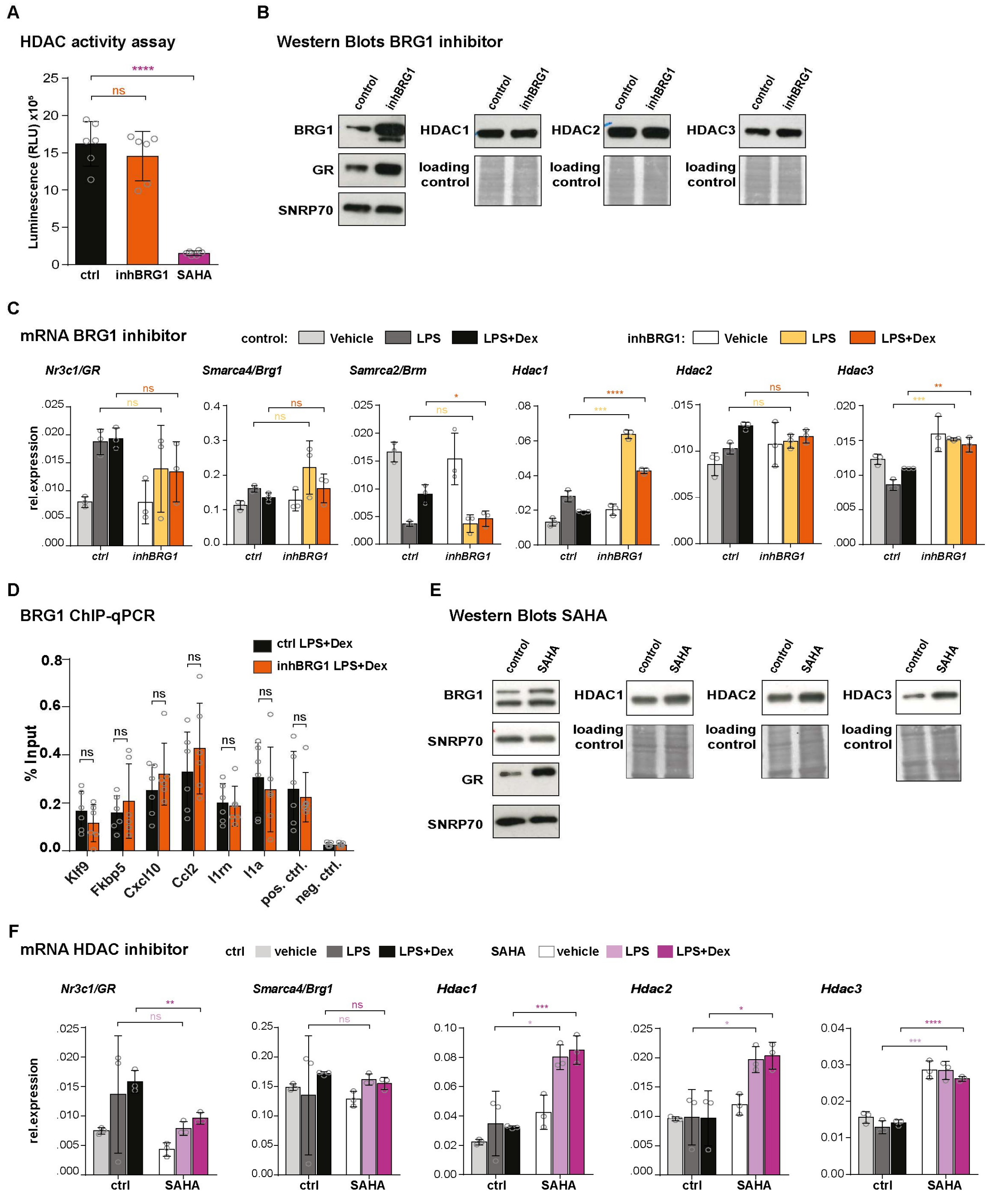


**Supplementary Figure 5. BRG1 and HDAC inhibition in primary macrophages** (A) HDAC activity assay in LPS+Dex stimulated macrophages treated with control, BRG1 inhibitor or SAHA. (B) Western blot showing nuclear BRG1, GR, HDAC1, HDAC2 and HDAC3 protein levels in control and BRG1 inhibitor treated macrophages (LPS+Dex). SNRP70 blotting and Naphthol Blue Black staining serve as loading control. (C) qRT-PCR of *GR, Brg1, Brm, Hdac1, Hdac2* and *Hdac3* in vehicle, LPS and LPS+Dex stimulated macrophages treated with vehicle (DMSO) or BRG1 inhibitor. (D) BRG1 ChIP-qPCR in LPS+Dex stimulated macrophages treated with vehicle (DMSO) or BRG1 inhibitor. (E) Western blot of nuclear BRG1, GR, HDAC1, HDAC2 and HDAC3 protein levels in control and SAHA treated macrophages (LPS+Dex). Loading controls: same as above. (F) qRT-PCR of *GR, Brg1, Brm, Hdac1, Hdac2* and *Hdac3* in vehicle, LPS and LPS+Dex stimulated macrophages treated with vehicle (DMSO) or SAHA. For all bar plots, values are mean ± standard deviation. $*p<0.05$, $**p<0.01$, $***p<0.001$, $****p<0.0001$, ns= not significant, unpaired two-tailed Student's t-test, n= 3.
